# Slide-level batch structure limits histology-guided supervision of transcriptomic foundation models

**DOI:** 10.64898/2026.09.30.755585

**Authors:** Ulysse McConnell, Kalin Nonchev, Viktor H Koelzer, Gunnar Rätsch

## Abstract

Spatial transcriptomics pairs spatially resolved gene expression with tissue morphology in the same tissue section. Transcriptomic foundation models encode such expression profiles into general-purpose representations, but these representations can retain slide- and cohort-specific variation that obscures biological signal. Here, we test whether matched H&E histology can improve these representations as a training-time supervisory signal that is discarded at inference. Across three transcriptomic foundation models, histology-guided supervision provides no consistent aggregate improvement in cross-donor annotation transfer, despite substantially stronger transfer from histology alone. We find that gene-expression embeddings from spot-based spatial transcriptomics are low-dimensional and strongly structured by slide identity, limiting the shared gene–morphology signal available for cross-modal transfer. Guidance benefits some morphologically distinctive classes but reduces held-out gene predictivity broadly across the transcriptome. These results suggest that cross-modal supervision can reorganize information already encoded in a frozen representation but cannot recover information the representation does not retain, highlighting the importance of diagnosing slide-specific structure before applying such supervision. Code availability: https://github.com/ratschlab/vision-guided-transcriptomics-fm-2026

## 1 Introduction

Spatial transcriptomics measures gene expression at known locations within a tissue section, often alongside imaging such as hematoxylin and eosin (H&E) histology. These data link molecular programs to tissue structure, with applications in disease classification [1–3] and tumor profiling [4– 6]. Each spatial location yields a high-dimensional gene-expression profile, motivating the use of transcriptomic foundation models to obtain compact representations for downstream analysis. These models are pre-trained with self-supervised objectives on large corpora of dissociated single-cell transcriptomes and provide general-purpose gene-expression representations. However, when applied to spatial transcriptomics data, their representations can retain noise and slide- or cohort-specific variation, limiting transfer to unseen donors [7].

In contrast, H&E morphology provides a well-established view of tissue organization [8], and pathology foundation models offer increasingly generalizable representations of this morphology [9– 11]. Because morphology is informative about molecular state [12], we hypothesize that these morphological representations can supervise a gene-expression encoder and make its features more transferable. Matched histology is often unavailable at deployment, so we provide it only during training and discard it at inference, a setting we term *histology-guided supervision*.

However, across three transcriptomic foundation models, histology-guided supervision provides no consistent aggregate improvement in cross-donor transfer, despite substantially stronger transfer from morphology alone. We find that the foundation-model representations are low-dimensional and strongly structured by slide-specific variation, leaving limited shared gene–morphology signal for transfer. Consistent with this limitation, histology guidance benefits some morphologically distinctive classes but can reduce held-out gene predictivity.

## 2 Related work

### 2.1 Foundation models

#### Pathology foundation models

These models use self-supervised objectives such as contrastive learning and masked image modeling to learn generalizable morphological representations from whole-slide images [9–11, 13]. Vision-language variants further align image tiles with text for classification with user-defined labels [14, 15].

#### Transcriptomic foundation models

These models use self-supervised objectives to learn generalizable gene-expression representations from large-scale single-cell and spatial transcriptomics data [16–19]. Beyond unimodal foundation models, multimodal models jointly learn from two or more modalities during pre-training, including combinations of histology, text, and omics [20–22]. In contrast, we use pre-trained histology and gene-expression models as separate, fixed feature extractors, isolating cross-modal supervision while avoiding multimodal inference.

### 2.2 Cross-modal learning

Using one modality as a teacher for another is well established [23] and can be viewed as *learning using privileged information* [24], where auxiliary information is available only during training. In spatial transcriptomics, prior work has predominantly used histology to predict gene expression, including with pathology foundation model features and multi-scale spatial context [25–28]. The reverse direction, using transcriptomic information to guide pathology models, has received less attention [29–31].

Recent work explores morphology-guided integration across tissue sections [32], distillation of microscopy features into gene-expression representations [33], joint spatial, transcriptomic and morphological representations [34], and cross-sample alignment for multimodal integration [35]. These approaches combine or align modalities, whereas we use morphology only as a training-time supervisory signal.

## 3 Methods

### 3.1 Histology-guided representation learning

Each Visium spot *i* has a frozen gene-expression embedding 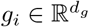and a morphology embedding 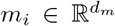. An encoder *f* maps *g*_*i*_ to *z*_*i*_ *∈* R^128^, and a decoder *ρ* predicts the corresponding morphology embedding (Figure 1):

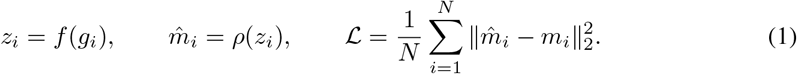

**Figure 1:**
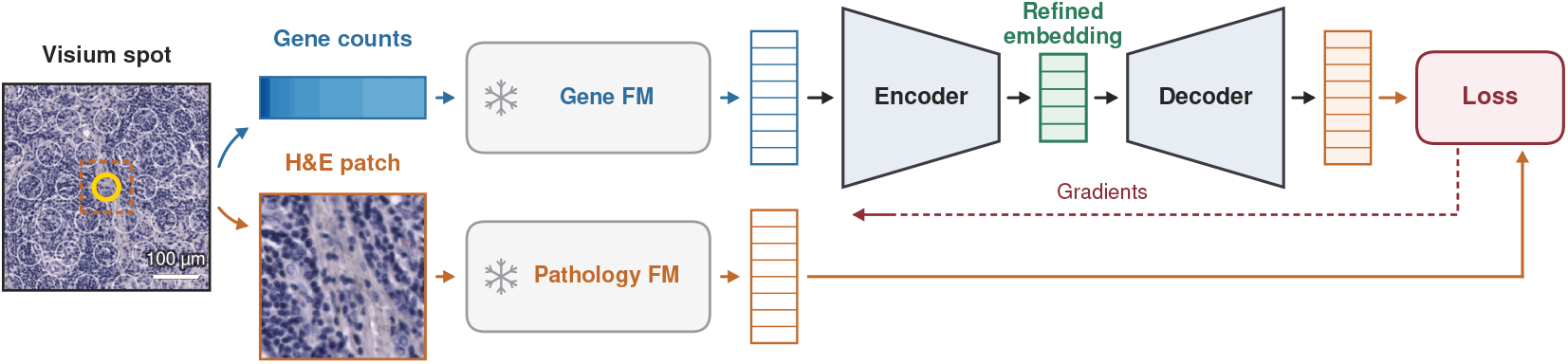
Histology-guided representation learning. Both foundation models are frozen. Morphology supervises the encoder through the reconstruction loss during training and is discarded at inference.

**Figure 2:**
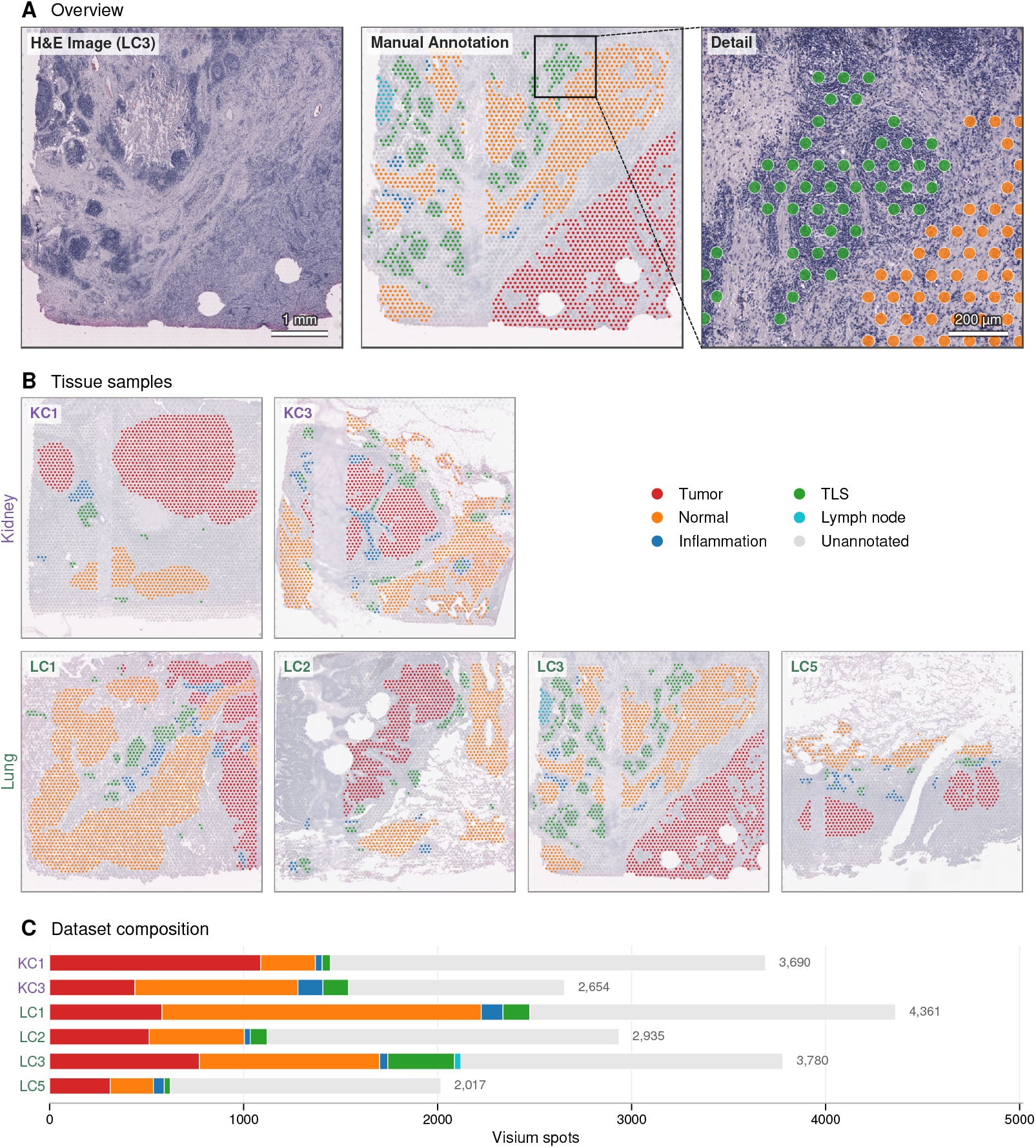
Overview of the USZ dataset. (A) Lung sample (LC3) with H&E image and overlaid expert annotation, (B) All six samples used for evaluation, (C) Composition of annotated tissue classes in each sample.

Morphology supervises *f* only through this reconstruction loss. At inference, *ρ* is discarded, and *z*_*i*_ depends only on gene expression. Architecture and training details are in Appendix C. Gene-expression embeddings are obtained from Geneformer V2 [17], scGPT [18], and CancerFoundation [36], and morphology embeddings from Midnight [13] (Appendix B), and only *f* and *ρ* are trained.

### 3.2 Data and evaluation

We train on 10x Visium spot-based data (55 *μ*m) from MOSAIC [37] and on lung samples from De Zuani et al. [38], and evaluate on USZ [39] (6 slides, 6 donors, kidney and lung) and Tumor Profiler (TuPro) [40, 34] (18 slides, 7 donors, metastatic melanoma) (Appendix A).

We evaluate frozen representations by leave-one-donor-out *k*-NN transfer (*k*=5) on expert annotations, measured by macro-F1, against PCA on gene-expression embeddings (128 components), a morphology-only PCA reference that reads H&E at inference, and a majority-class baseline (Appendix D). The probe is fitted on the remaining donors, which prior work terms zero-shot [7]. Gene selection for scGPT and CancerFoundation is fitted once over the pooled cohort including held-out slides, so preprocessing is transductive although it uses no annotations (Appendix B). We characterize the frozen embeddings using effective rank [41], the leading gene–morphology canonical correlation and the fraction of it accounted for by slide identity, and iLISI [42]. To assess the role of spot-level correspondence, we repeat training after shuffling H&E patches within slides or globally across spots.

## 4 Results

Table 1 summarizes cross-donor annotation transfer on TuPro for the three foundation models. Gene-only PCA achieves macro-F1 scores of 0.53–0.55, whereas histology-guided representations achieve 0.52–0.56, within 0.02 of the gene-only baseline for every model. The frozen embeddings themselves offer little advantage over HVG + PCA on raw counts, echoing reported zero-shot limitations of single-cell foundation models [7]. The morphology-only PCA reference reaches 0.77, so morphology contains far more transferable information than guidance recovers. Per-organ results show the same pattern, except that guidance improves a weak Geneformer V2 kidney baseline (Appendix E.1).

**Table 1:** Cross-donor *k*-NN annotation transfer macro-F1 (*k*=5) on TuPro skin melanoma (7 donors). Brackets give the 95% CI from a donor bootstrap, AE is the morphology-reconstruction encoder, and CDANN is the control variant per foundation model.

|  | PCA (gene-only) | AE (gene→H&E) | CDANN |
| --- | --- | --- | --- |
| HVG + PCA (counts) |  | 0.54 [0.46, 0.60] |  |
| PCA reference (H&E) |  | 0.77 [0.72, 0.82] |  |
| *Transcriptomic foundation model + Midnight H&E* |  |  |  |
| CancerFoundation | 0.55 [0.48, 0.59] | 0.54 [0.47, 0.61] | 0.56 [0.50, 0.61] |
| scGPT | 0.53 [0.47, 0.59] | 0.52 [0.44, 0.58] | 0.53 [0.46, 0.62] |
| Geneformer V2 | 0.54 [0.47, 0.60] | 0.52 [0.45, 0.59] | 0.56 [0.49, 0.62] |
| Majority baseline |  | 0.20 [0.18, 0.22] |  |

### 4.1 Gene-expression embeddings are low-dimensional and strongly structured by slide

The frozen gene-expression embeddings have effective rank below 30 (28.9, 27.1, and 17.1 for Geneformer V2, scGPT, and CancerFoundation, in 1152, 512, and 256 dimensions), so most of their ariation lies in a small subspace, whereas the Midnight morphology embeddings have effective rank 80.5 in 3072 dimensions. Low rank does not by itself imply poor biological content, as integration methods such as sysVI operate at effective rank 8.4 in 50 dimensions [43]. It matters here because the subspace is strongly aligned with slide identity.

Slide identity also dominates the cross-modal association. Although the leading canonical correlation between gene expression and morphology is close to 1 in-sample, about 90% of it is accounted for by slide identity. iLISI likewise indicates poor cross-slide mixing for all three embeddings, so the shared signal available for transfer is confounded by slide-level structure (Appendix F). Because slide identity can reflect technical as well as biological differences, we interpret this as confounding by slide-specific variation rather than direct evidence of technical batch effects.

Post-embedding correction with Harmony [42] or ComBat [44] does not consistently improve transfer and requires joint data access, so we report it as a transductive diagnostic (Appendix E.2) [7].

### 4.2 Morphology provides class-specific benefits

The effect of guidance varies substantially across tissue classes (Table 2), with the morphologically distinctive tumor classes in USZ responding most clearly. Histology guidance increases tumor F1 from 0.46 to 0.53 in kidney and from 0.52 to 0.59 in lung. This benefit is lost under global shuffling, and in kidney also under within-slide shuffling. In lung, however, matched and within-slide guidance perform similarly, suggesting a tissue-dependent role of spot-level correspondence. Classes with less distinctive morphology show little change or a decrease, most clearly tertiary lymphoid structures.

**Table 2:** Per-class cross-donor *k*-NN annotation F1 (*k*=5, scGPT). *Matched* uses true gene–patch pairs, shuffles permute them within slides or globally, and brackets give 95% donor-bootstrap CIs.

| Class (support) | PCA | AE (gene→H&E) |  |  |
| --- | --- | --- | --- | --- |
|  | (gene-only) | Matched | Within-slide shuffle | Global shuffle |
| *TuPro*, skin melanoma (18 slides, 7 donors) |  |  |  |  |
| Tumor (52,240) | 0.86 [0.77, 0.93] | 0.86 [0.77, 0.92] | 0.86 [0.78, 0.92] | 0.85 [0.76, 0.91] |
| Normal lymphoid (12,938) | 0.67 [0.46, 0.78] | 0.66 [0.44, 0.78] | 0.63 [0.42, 0.75] | 0.65 [0.44, 0.75] |
| Stroma (9,969) | 0.54 [0.36, 0.73] | 0.50 [0.34, 0.67] | 0.46 [0.33, 0.62] | 0.48 [0.31, 0.63] |
| *USZ*, kidney carcinoma (2 slides, 2 donors) |  |  |  |  |
| Tumor (1,528) | 0.46 [0.39, 0.50] | 0.53 [0.43, 0.63] | 0.33 [0.11, 0.56] | 0.32 [0.03, 0.53] |
| Tertiary lymphoid structures (174) | 0.51 [0.46, 0.52] | 0.40 [0.33, 0.43] | 0.14 [0.03, 0.19] | 0.48 [0.18, 0.62] |
| Normal (1,122) | 0.48 [0.31, 0.61] | 0.45 [0.38, 0.57] | 0.44 [0.33, 0.56] | 0.38 [0.31, 0.50] |
| *USZ*, lung carcinoma (4 slides, 4 donors) |  |  |  |  |
| Tumor (2,177) | 0.52 [0.23, 0.66] | 0.59 [0.35, 0.71] | 0.59 [0.41, 0.69] | 0.46 [0.18, 0.60] |
| Tertiary lymphoid structures (601) | 0.52 [0.42, 0.55] | 0.33 [0.25, 0.48] | 0.24 [0.12, 0.31] | 0.33 [0.27, 0.45] |
| Normal (3,288) | 0.65 [0.38, 0.75] | 0.64 [0.57, 0.69] | 0.59 [0.45, 0.64] | 0.59 [0.35, 0.66] |

### 4.3 Histology-guided supervision can reduce molecular predictivity

We next ask whether guidance preserves the information in the original embeddings, using held-out per-gene ridge *R*^2^ (Appendix E.3). Guidance reduces this *R*^2^ for all three models. Across donors, it falls by 49% for CancerFoundation (mean Δ*R*^2^ = *−* 0.032), 46% for Geneformer V2, and 28% for scGPT, with smaller within-donor decreases. The loss is largely independent of baseline predictability, and no Hallmark program is consistently spared, so guidance reduces held-out predictivity broadly, not selectively.

## 5 Discussion & Conclusion

In this work, we tested whether matched H&E histology can act as privileged supervision for transcriptomic foundation models, training a gene-expression encoder that operates without histology at inference. Morphology is a biologically motivated teacher, and the morphology-only reference transfers better than any gene-based representation we tested, so we expected guidance to help.

Our results show no consistent aggregate improvement in cross-donor transfer for any of the three foundation models. The frozen embeddings occupy fewer than 30 effective dimensions, and much of their variation shared with morphology is associated with slide identity rather than transferable biological variation. Accordingly, guidance benefits morphologically distinctive classes but reduces held-out gene predictivity across the transcriptome. This is consistent with frozen-encoder fusion exploiting shared rather than synergistic information [45].

In summary, cross-modal supervision can reorganize information already present in a frozen representation, but cannot recover information it does not retain. Source-embedding dimensionality and slide-specific structure should therefore be diagnosed before applying privileged supervision, and such guidance should be evaluated for the information it removes, not only downstream accuracy.

### Limitations

Our study uses a single pathology foundation model and two held-out cohorts. The evaluation mixes donor, cohort, and slide shifts, preventing isolation of donor effects. Morphology patches use a fixed pixel size without stain normalization, which may introduce slide-specific structure. Finally, the transcriptomic models are pre-trained on dissociated single cells rather than multi-cell Visium spots, which may depress effective rank.

#### Use of large language models

Generative AI tools, including large language models such as OpenAI ChatGPT and Anthropic Claude and code-completion assistants such as GitHub Copilot, were used during manuscript preparation to assist with prose editing, literature organization, and software development. All AI-generated content was critically reviewed, edited, and verified by the authors, who take full responsibility for the accuracy, integrity, and originality of the final manuscript. No AI tool was used to generate scientific ideas, interpret experimental results, or draw scientific conclusions.

## Acknowledgments and Disclosure of Funding

Computational data analysis was performed at Leonhard Med (https://sis.id.ethz.ch/services/sensitiveresearchdata/), a secure trusted research environment at ETH Zurich.

This study makes use of data generated by the MOSAIC consortium (Owkin, Charité – Universitätsmedizin Berlin, Lausanne University Hospital - CHUV, Erlangen Hospital, Gustave Roussy Institute, University of Pittsburgh) and made available through the MOSAIC Window initiative. Readers should note that the MOSAIC consortium bears no responsibility for the further analysis or interpretation of these data beyond what is published by the MOSAIC consortium partners.

We also use data from the Tumor Profiler (TuPro) study [46] and thank the Tumor Profiler Consortium (https://tumorprofilercenter.ch).

## A Datasets

We train on Visium spatial transcriptomics data from MOSAIC [37] (covering samples from glioblastoma, diffuse large B-cell lymphoma, bladder cancer, mesothelioma, and ovarian cancer) together with two lung datasets covering lung adenocarcinoma (LUAD) and lung squamous cell carcinoma (LUSC) [38]. For evaluation, we use two datasets from USZ [39] (covering kidney and lung cancer) and a skin cancer dataset from the Tumor Profiler study [40, 34]. The USZ and Tumor Profiler datasets additionally contain annotations of tissue types, allowing us to compute classification scores on these slides.

Table 3 summarizes the composition of all datasets, and Supplementary Figure 2 shows the annotated tissue classes of the USZ evaluation slides.

**Table 3:**
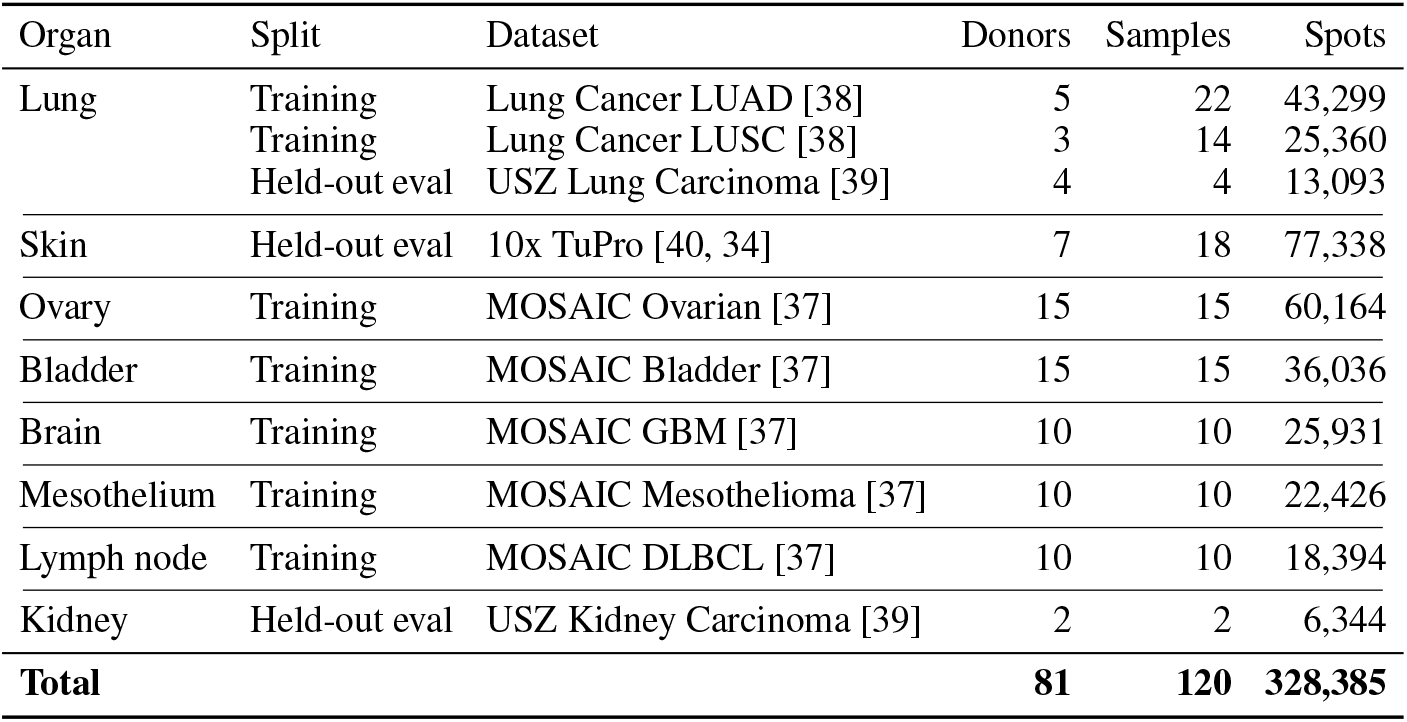
Dataset composition.

**Table 4:** Frozen foundation model checkpoints. Revisions are the leading eight characters of the commit hash.

| Model | Revision | Source |
| --- | --- | --- |
| Geneformer V2 | 04c2b2e8 | https://huggingface.co/ctheodoris/Geneformer |
| CancerFoundation | 130375b3 | https://github.com/BoevaLab/CancerFoundation |
| scGPT (scGPT_human) | † | https://github.com/bowang-lab/scGPT |
| Midnight | adc6b156 | https://huggingface.co/kaiko-ai/midnight |

† scGPT distributes scGPT\_human as an unversioned file download, and the checkpoint used here has md5 9922ec94305126e6e4f9c1575cf493ae.

**Table 5:** Fold hierarchy. *Held out in* lists the levels at which a sample is held out, namely replicate (rep), region (reg), and donor.

| Organ | Donor | Region | Sample ID | Held out in |
| --- | --- | --- | --- | --- |
| Kidney | KC1 | – | KC1 | donor |
|  | KC3 | – | KC3 | donor |
| Lung | LC1 | – | LC1 | donor |
|  | LC2 | – | LC2 | donor |
|  | LC3 | – | LC3 | donor |
|  | LC5 | – | LC5 | donor |
| Skin | MACEGEJ | Region 1 | MACEGEJ-1-1 | rep, reg, donor |
|  |  |  | MACEGEJ-1-2 | rep, reg, donor |
|  |  | Region 2 | MACEGEJ-2-1 | rep, reg, donor |
|  |  |  | MACEGEJ-2-2 | rep, reg, donor |
|  | MAHEFOG | Region 1 | MAHEFOG-1-1 | rep, donor |
|  |  |  | MAHEFOG-1-2 | rep, donor |
|  | MAJOFIJ | Region 1 | MAJOFIJ-1-1 | rep, reg, donor |
|  |  |  | MAJOFIJ-1-2 | rep, reg, donor |
|  |  | Region 2 | MAJOFIJ-2-1 | rep, reg, donor |
|  |  |  | MAJOFIJ-2-2 | rep, reg, donor |
|  | MAKYGIW | Region 1 | MAKYGIW-1-1 | rep, donor |
|  |  |  | MAKYGIW-1-2 | rep, donor |
|  | MANOFYB | Region 1 | MANOFYB-1-1 | rep, donor |
|  |  |  | MANOFYB-1-2 | rep, donor |
|  | MELIPIT | Region 1 | MELIPIT-1-1 | rep, donor |
|  |  |  | MELIPIT-1-2 | rep, donor |
|  | MIDEKOG | Region 1 | MIDEKOG-1-1 | rep, donor |
|  |  |  | MIDEKOG-1-2 | rep, donor |

## B Feature extraction

Gene-expression embeddings are computed per spot from raw counts. Geneformer V2 uses the repository-root checkpoint (18 layers, width 1152, 4096-token context) with Ensembl identifiers and no gene selection, and the embedding is the [CLS] token of the second-to-last layer. scGPT uses the whole-human checkpoint scGPT_human after normalization to 10^4^ counts per spot, log1p, and selection of 2000 highly variable genes (seurat_v3, cohort as batch key). CancerFoundation receives raw counts and selects 1199 highly variable genes (cell_ranger) internally. Both use a 1200-token context and return the *ℓ*_2_-normalized [CLS] token, and for both the selection is made once over the pooled cohort so that all slides share one gene set. The selection uses no annotation labels but sees the held-out slides, so preprocessing for these two models is transductive rather than strictly inductive. Genes are retained if detected in at least 100 spots and non-zero in every cohort. No spot-level filtering is applied, as a depth threshold would preferentially remove tissue-border spots. The resulting dimensions are 1152, 512, and 256.

Morphology embeddings are computed with Midnight from one 224 *×* 224 patch per spot, cut at the spot center in the full-resolution image and not rescaled, with the model’s own channel normalization and no stain normalization. The embedding concatenates the [CLS] token with the mean of the patch tokens (3072 dimensions). Morphology embeddings enter the reconstruction loss unstandardized and unwhitened.

## C Training methodology

Encoder *f* and decoder *ρ* are MLPs of three linear layers, with hidden widths interpolated geometrically between their input and output dimensions (e.g. on Geneformer V2 the encoder is 1152 *→* 512 *→* 256 *→* 128 and the decoder 128 *→* 512 *→* 1024 *→* 3072). Hidden layers use batch normalization, GELU, and dropout 0.1, and the output layer of each is left bare. A LayerNorm on the encoder output is part of the deployed embedding, *z*_*i*_ = LayerNorm(*f* (*g*_*i*_)), and fixes the latent scale so that *k*-NN distances are comparable across representations. Training uses AdamW [47] with a learning rate of 10^*−*3^ under cosine annealing, weight decay 10^*−*2^, and batch size 4096, for up to 200 epochs. A random 10% of the training spots is held out for early stopping (patience 20) and the best-validation weights are restored.

### C.1 CDANN model

The CDANN model is a conditional domain-adversarial neural network, which asks whether guidance improves once slide identity is penalized directly instead of being left to the reconstruction. Projectors *f* and *q* map the gene and morphology embeddings to R^128^ through one hidden layer of width 512 with LayerNorm, GELU, and dropout 0.1. Over a batch of *N* spots,

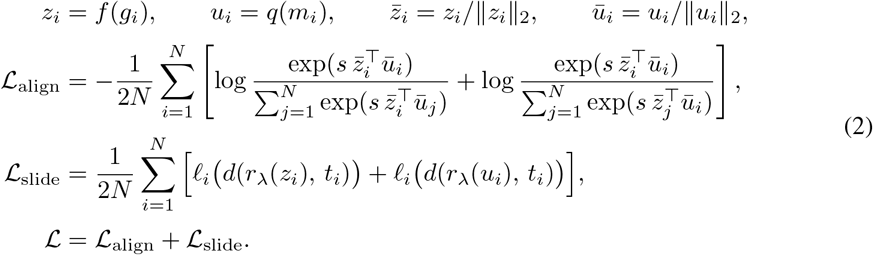

ℒ_align_ is a symmetric InfoNCE loss over the batch, in which the matched spot is the positive pair and the other *N −* 1 spots are negatives, and the temperature *s* is learned and initialized at 1*/*0.07 [48]. In ℒ_slide_, a discriminator *d* predicts the slide of spot *i* from each projection concatenated with the one-hot tissue *t*_*i*_, and *ℓ*_*i*_ is the cross-entropy against the true slide with label smoothing 0.1, its softmax and smoothing mass restricted to the slides of that tissue, so the discriminator separates slides within a tissue rather than tissues. The gradient reversal *r*_*λ*_ [49] is the identity in the forward pass and scales the gradient by *− λ* in the backward pass, so one descent step on ℒ trains *d* to recover the slide and *f, q* to hide it, and *− λ* is ramped from 0 to 0.5 over training. Projectors and discriminator have separate AdamW optimizers (learning rates 10^*−*4^ and 10^*−*5^, weight decay 10^*−*2^, batch size 256) and train for a fixed 50 epochs, and only the gene projector is deployed.

## D Evaluation methodology

We give an overview of the hierarchical structure of the evaluation dataset in Supplementary Table 5. Predictive performance on the pathology annotation classes is evaluated leave-one-donor-out (LODO) within each tissue. For each donor in turn, a *k*-NN classifier (*k*=5) is fitted on the annotated spots of that tissue’s remaining donors and applied to the held-out donor’s spots. The probe’s training set is capped at 10,000 spots by class-stratified subsampling, and embeddings are standardized using the mean and variance of those training spots alone, while held-out spots are not subsampled. This procedure yields one prediction per annotated spot by a classifier that never saw that spot’s donor.

Predictions are obtained across three training seeds (42, 43, 44). For a given tissue and training seed, confidence intervals are obtained by a donor-level bootstrap over the fixed predictions. We repeatedly draw a donor with replacement, pool all spots of the drawn donors with multiplicity, and recompute the macro-F1 from the resulting confusion matrix.

A pathology annotation class is scored only if it is supported by at least 50 annotated spots on at least 2 slides *within its own tissue*, effectively dropping the lung lymph node and skin pigment classes.

## E Extended results

### E.1 Cross-donor annotation transfer

Each representation is fitted once per seed on the 231,610 spots of the 96 unannotated training slides and then applied unchanged to all 86,614 annotated spots. Transfer is then scored with the leave-one-donor-out probe of Appendix D, run within organ, giving 13 folds over kidney (2 donors, 2,988 annotated spots), lung (4 donors, 6,306 annotated spots), and skin (7 donors, 77,320 annotated spots).

An organ’s fold predictions are pooled and scored once, so a macro-F1 is one average over the union of held-out donors rather than a mean of per-fold scores. The average runs over all scored classes in every organ. Point estimates are the mean over the three training seeds. Each seed contributes 2000 donor-bootstrap replicates, and the reported interval is the 2.5th to 97.5th percentile of the three sets taken together, so it covers donor sampling and initialization.

Table 6 compares six approaches, with every learned representation at width 128. *HVG + PCA* applies the standard scanpy workflow to raw counts, namely CP10k, log1p, 2000 highly variable genes ranked by seurat_v3, per-gene scaling, and PCA, with the ranking and the scaler fitted on the training split alone. *PCA (gene-only)* is PCA of the frozen gene-expression embedding. *AE* is the morphology-reconstruction encoder of Section 3. *CDANN* is a conditional domain-adversarial encoder described in Appendix C.1. The *PCA reference* is PCA of the frozen Midnight embedding and reads morphology at inference, so it is not a deployable baseline. The *majority* rule predicts the most frequent training class.

**Table 6:** Cross-donor *k*-NN annotation transfer macro-F1 (*k*=5) per organ, for every foundation model. Brackets give the 95% CI from a donor bootstrap

| Model | Kidney (2) | Lung (4) | Skin (7) |
| --- | --- | --- | --- |
| *Substrate-independent baselines* |  |  |  |
| HVG + PCA (counts) | 0.35 [0.19, 0.47] | 0.35 [0.23, 0.49] | 0.53 [0.47, 0.59] |
| PCA reference (H&E) | 0.59 [0.46, 0.79] | 0.79 [0.69, 0.83] | 0.77 [0.72, 0.82] |
| Majority baseline | 0.13 [0.08, 0.13] | 0.17 [0.14, 0.19] | 0.20 [0.18, 0.22] |
| *CancerFoundation + Midnight H&E* |  |  |  |
| PCA (gene-only) | 0.45 [0.44, 0.48] | 0.43 [0.28, 0.51] | 0.55 [0.48, 0.59] |
| AE (gene→H&E) | 0.46 [0.37, 0.52] | 0.44 [0.33, 0.51] | 0.54 [0.47, 0.61] |
| CDANN | 0.46 [0.42, 0.60] | 0.39 [0.28, 0.49] | 0.56 [0.50, 0.61] |
| *scGPT + Midnight H&E* |  |  |  |
| PCA (gene-only) | 0.37 [0.33, 0.38] | 0.44 [0.33, 0.48] | 0.53 [0.47, 0.59] |
| AE (gene→H&E) | 0.35 [0.29, 0.41] | 0.40 [0.32, 0.46] | 0.52 [0.44, 0.58] |
| CDANN | 0.38 [0.35, 0.47] | 0.36 [0.26, 0.46] | 0.53 [0.46, 0.62] |
| *Geneformer V2 + Midnight H&E* |  |  |  |
| PCA (gene-only) | 0.19 [0.18, 0.29] | 0.49 [0.41, 0.55] | 0.54 [0.47, 0.60] |
| AE (gene→H&E) | 0.46 [0.28, 0.54] | 0.45 [0.36, 0.53] | 0.52 [0.45, 0.59] |
| CDANN | 0.45 [0.39, 0.58] | 0.46 [0.37, 0.54] | 0.56 [0.49, 0.62] |

**Table 7:**
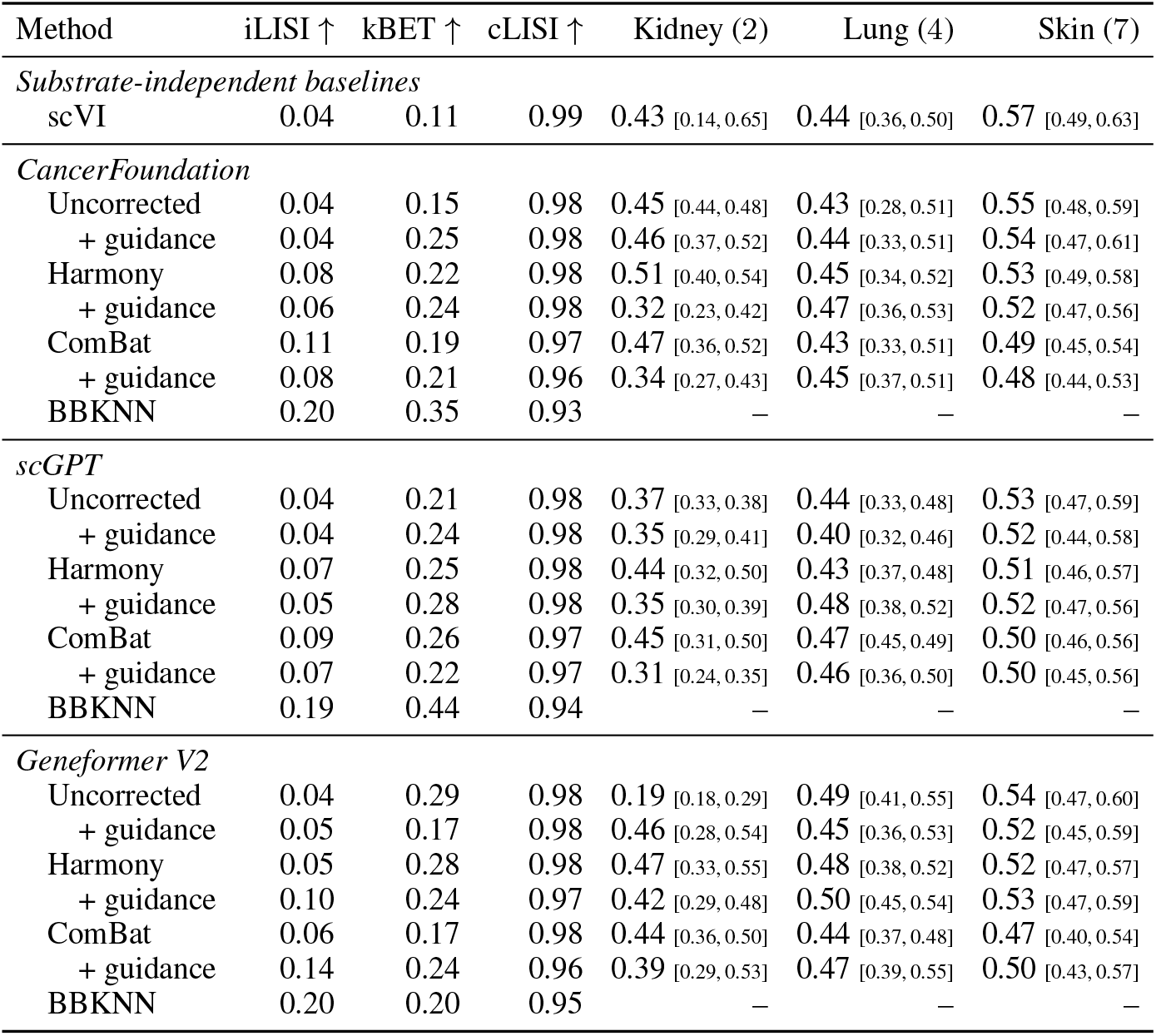
Batch correction applied to the frozen embeddings, with the probe reported per tissue. iLISI and kBET measure batch mixing, while cLISI measures conservation of biological label structure. Brackets give the 95% CI from a donor bootstrap within each tissue.

### E.2 Batch correction

We apply different batch-correction methods to the gene-expression embeddings, including Harmony [42], BBKNN [50], and ComBat [44] to test the impact on downstream guidance. Batch mixing is quantified in terms of integration LISI (iLISI) [42] and kBET [51], while preservation of biological structure is measured in terms of cell-type LISI (cLISI). Each correction is fitted *once and globally* (including the evaluation data), with the slide as the batch covariate. Because these methods require joint access to all batches, we report them as transductive diagnostics rather than as transfer baselines. The corrected embeddings are then used in the histology-guided representation learning as new frozen gene-expression embeddings. In addition, we use scVI [52] on the raw gene expression with the batch as covariate to produce embeddings. The *uncorrected* rows correspond to the setting described in Section 3. We note that BBKNN corrects the neighbor graph rather than the embeddings, so no separate F1 scores are reported. Post-embedding correction improves slide mixing only marginally, iLISI barely increases and kBET stays below 0.30, and does not consistently improve downstream macro-F1.

### E.3 Per-gene predictivity

Supplementary Figure 3 accompanies Section 4.3 and compares per-gene *R*^2^ for the frozen Cancer-Foundation embeddings against the guided embeddings, both across replicates and across donors.

Targets are the 15,944 genes shared by every annotated slide and detected in at least 10 of 40,000 subsampled spots, normalized to 10^4^ counts per spot and log(1 + *x*) transformed. Targets and the representation being scored are standardized on each fold’s training spots, so no intercept is fitted. The ridge penalty is selected per gene, fold, and representation by generalized cross-validation over 33 log-spaced values in [10^*−*1^, 10^7^] using training rows only, and each fold is capped at 30,000 training spots. Both levels are constructed within the TuPro skin cohort (Table 5). Cross-replicate (18 folds) holds out one replicate, and cross-donor (7 folds) holds out one donor. Held-out predictions are pooled over a level’s folds and scored per gene against the pooled held-out mean. The negative values in Figure 3 are shallow, at medians of *−* 0.007 across replicates and *−* 0.012 across donors. A gene below zero is one the embedding does not predict rather than one it predicts badly.

**Figure 3:**
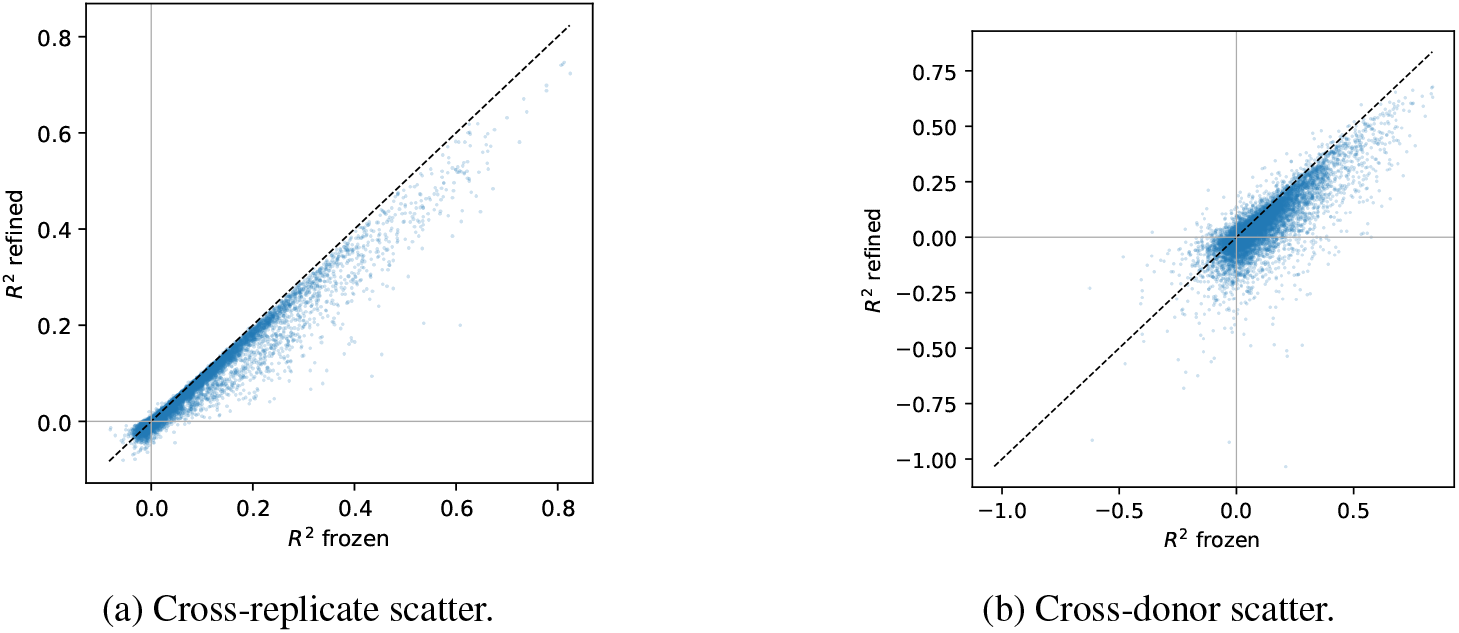
Per-gene *R*^2^ for the frozen CancerFoundation embeddings against the guided embeddings from the encoder-decoder, under cross-replicate and cross-donor held-out settings.

**Figure 4:**
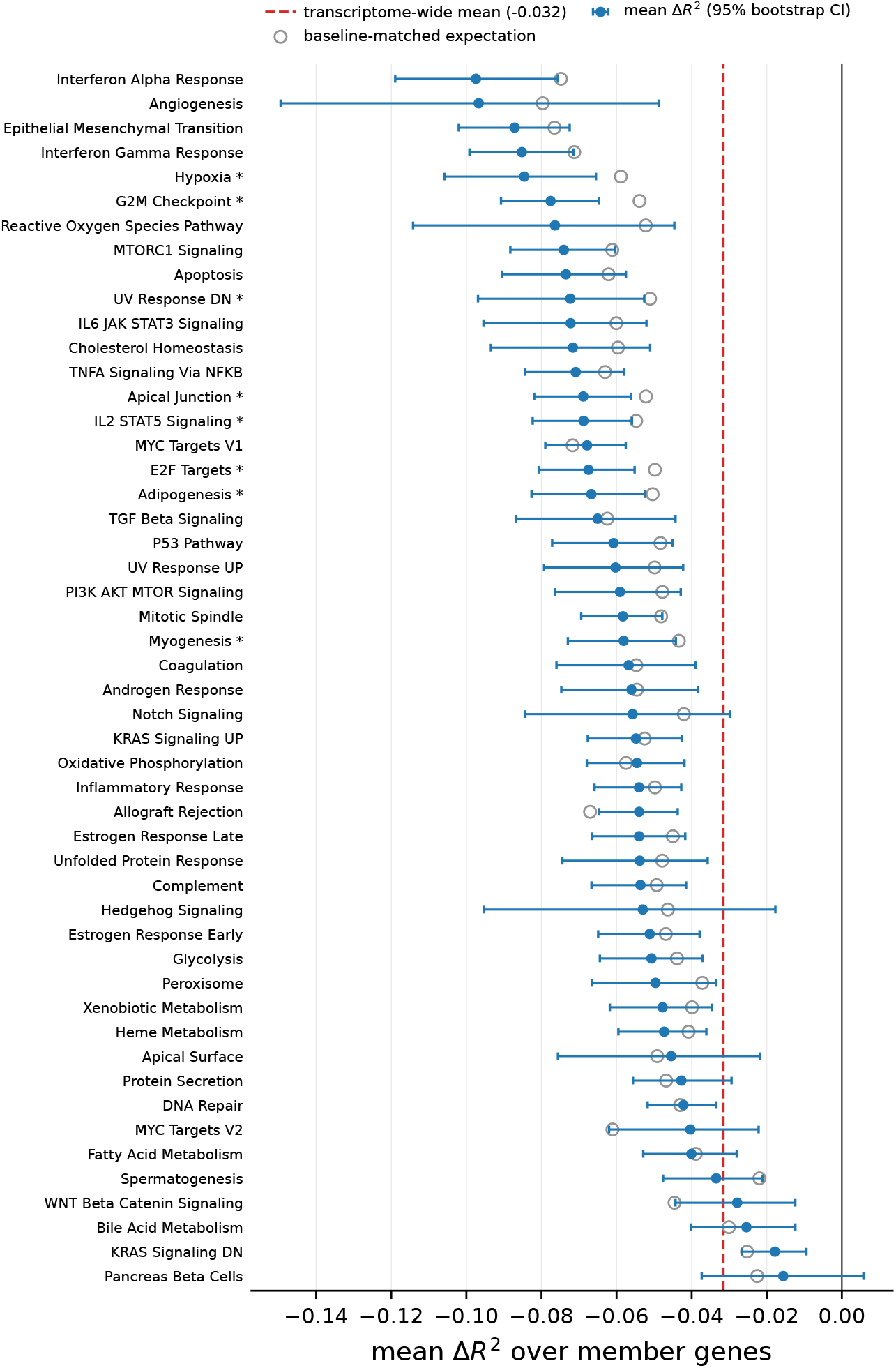
The 50 MSigDB Hallmark sets ranked by mean Δ*R*^2^ over member genes, frozen against guided CancerFoundation embeddings across donors. Bars are 95% percentile bootstrap intervals, and the dashed line is the transcriptome-wide mean Δ*R*^2^ of *−* 0.032. Open markers give each set’s expectation under the baseline-matched permutation null (20 equal-count bins of the frozen *R*^2^); stars mark sets displaced from it at *q <* 0.05.

**Figure 5:**
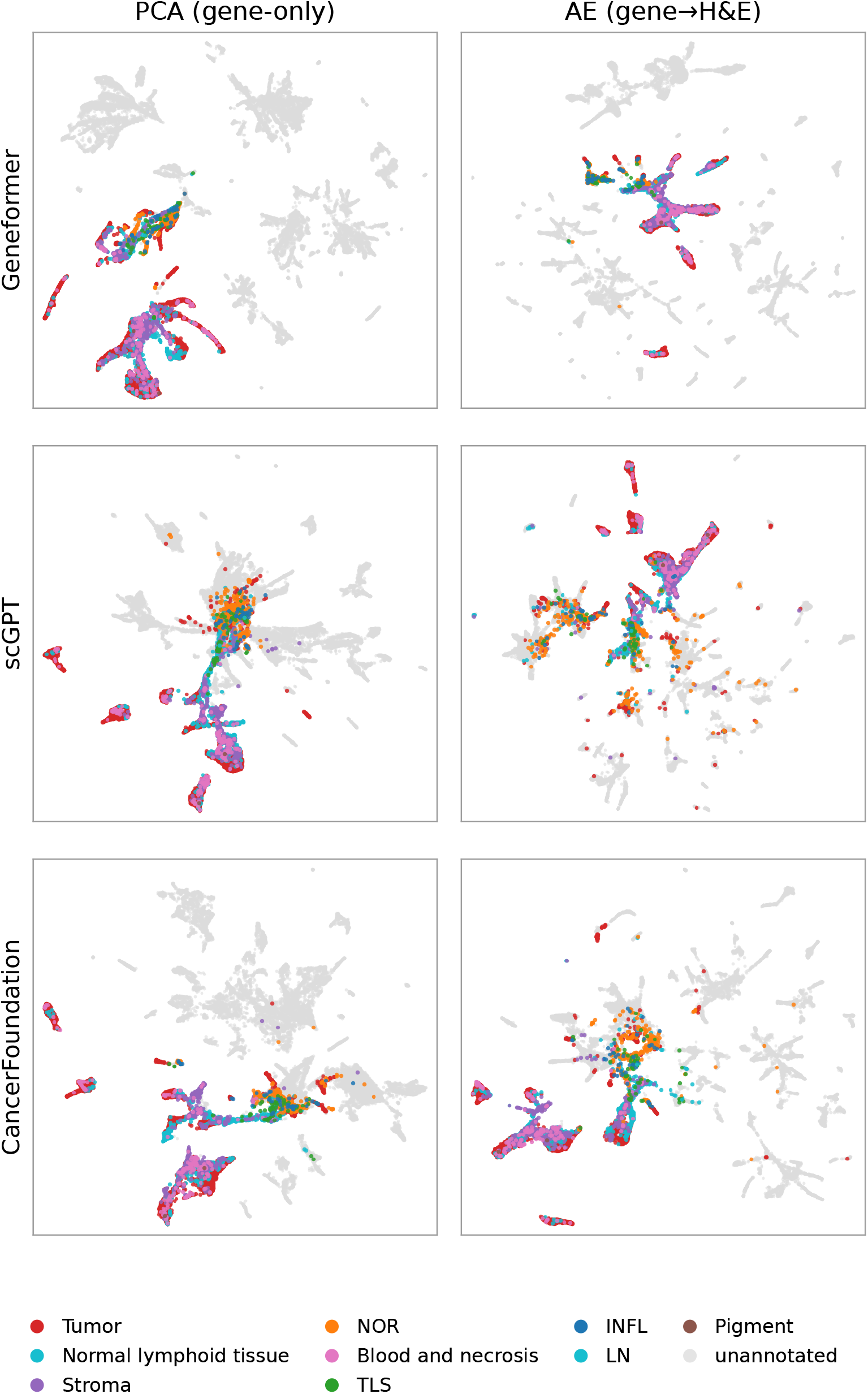
UMAP of gene-expression embeddings across all datasets colored by annotation where available.

**Figure 6:**
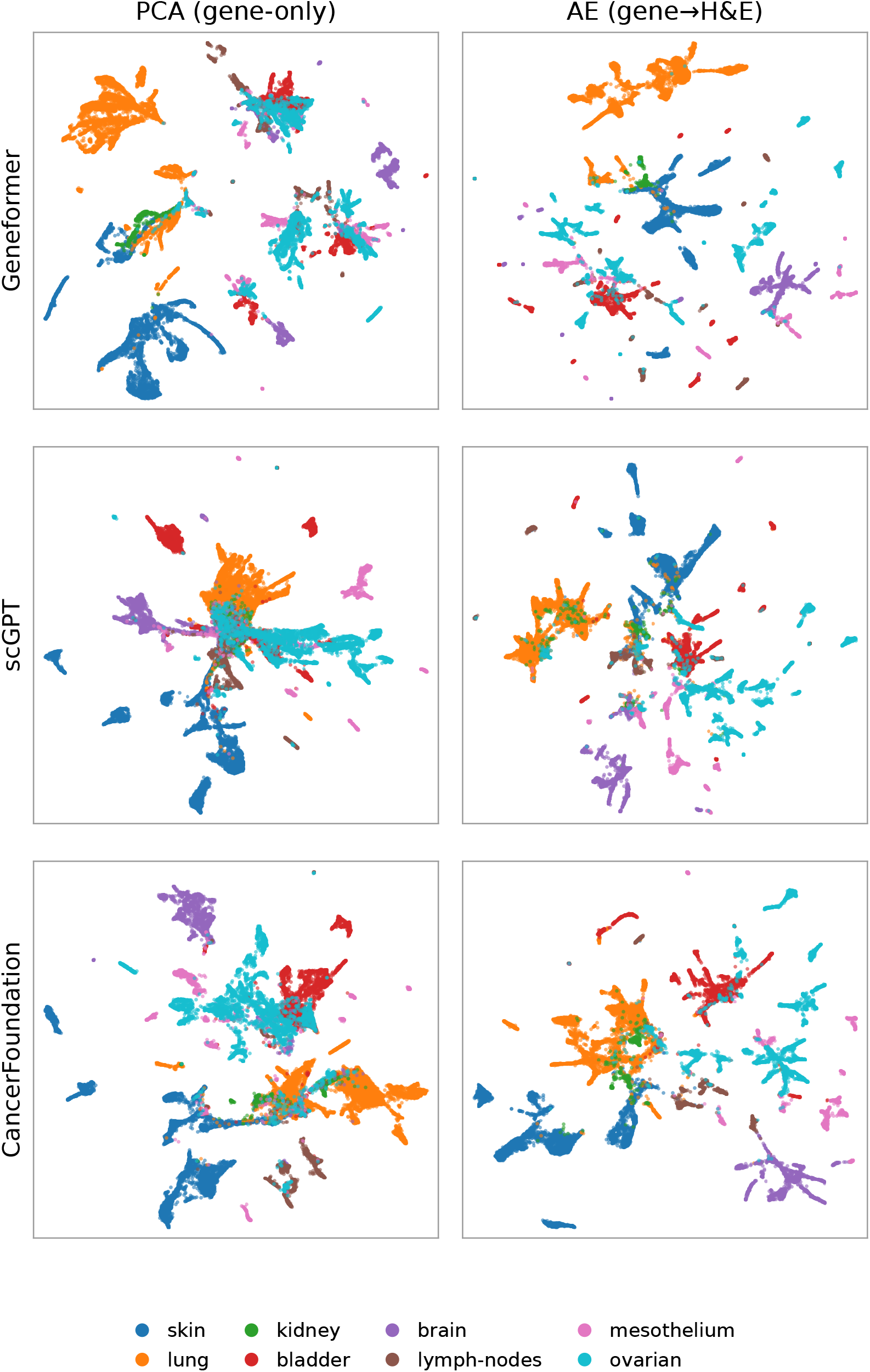
UMAP of gene-expression embeddings across all datasets colored by tissue type.

**Figure 7:**
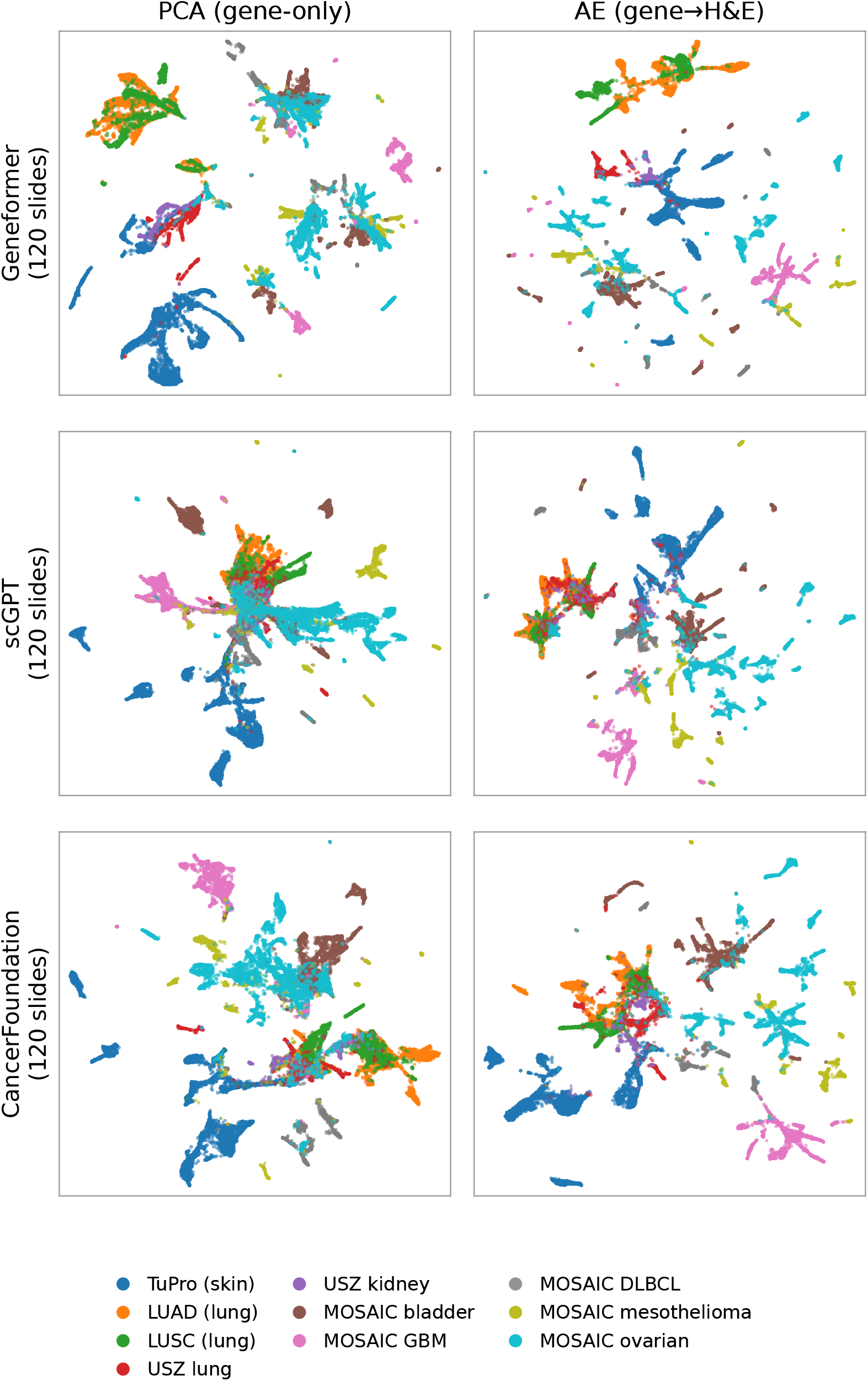
UMAP of gene-expression embeddings across all datasets colored by dataset.

Gene sets are MSigDB Hallmark [53] (50 sets, 6,971 HGNC symbols) retrieved with decoupler [54].^6^ Symbols are matched directly with no identifier mapping, 3,957 are present in the 15,944-gene background, and all 50 sets exceed the minimum of 15 members. A set is summarized by the mean Δ*R*^2^ over its members and compared against the transcriptome-wide mean by two-sided permutation over 10,000 gene-label shuffles, with Benjamini–Hochberg correction. Guidance costs predictivity in proportion to how predictable a gene was to begin with (*ρ* = *−* 0.80 between a set’s mean Δ*R*^2^ and its mean frozen *R*^2^), and curated sets are built from well-predicted genes, so the shuffle is repeated within 20 equal-count bins of the frozen *R*^2^. This second null asks whether a set is displaced beyond what its composition already implies.

Across donors on CancerFoundation every Hallmark set moves down, but against the baseline-matched null eight sets are displaced beyond their composition (Hypoxia, G2M Checkpoint, UV Response DN, E2F Targets, Apical Junction, Adipogenesis, Myogenesis, and IL2/STAT5 Signaling), and between 7 and 19 sets over the three foundation models and both levels. Guidance costs predictivity broadly rather than in specific pathways.

## F Visualization of embeddings

Supplementary Figures 5 to 7 illustrate the slide- and dataset-specific structure in the gene-expression embeddings through UMAP plots, colored by annotation, tissue type, and dataset, respectively.

## Footnotes

6 dc.op.hallmark does not stamp an MSigDB release. Every table here was scored against one vendored copy, retrieved with decoupler 2.1.6, which has md5 7ee7c3a1b484cc86d69caa94268dfe5c.

## Notes

https://github.com/ratschlab/vision-guided-transcriptomics-fm-2026

### Competing Interest Statement

The authors have declared no competing interest.

